# Maternal Obesity Enhances Oocyte Chromosome Abnormalities Associated With Aging

**DOI:** 10.1101/505511

**Authors:** Yan Yun, Zijie Wei, Neil Hunter

## Abstract

Obesity is increasing globally and maternal obesity has adverse effects on pregnancy outcomes and the long-term health of offspring. Maternal obesity has been associated with pregnancy failure through impaired oogenesis and embryogenesis. However, whether maternal obesity causes chromosome abnormalities in oocytes has remained unclear. Here we show that chromosome abnormalities are increased in the oocytes of obese mice and identify weakened sister-chromatid cohesion as the likely cause. Numbers of full-grown follicles retrieved from obese mice were the same as controls and the efficiency of *in vitro* oocyte maturation remained high. However, chromosome abnormalities presenting in both metaphase-I and metaphase-II were elevated, most prominently the premature separation of sister chromatids. Weakened sister-chromatid cohesion in oocytes from obese mice was manifested both as the terminalization of chiasmata in metaphase-I and as increased separation of sister centromeres in metaphase II. Obesity-associated abnormalities were elevated in older mice implying that maternal obesity exacerbates the deterioration of cohesion seen with advancing age.

## INTRODUCTION

Aneuploid embryos are a major cause of human infertility, miscarriage and congenital disease (Herbert et al. 2015; Jones and Lane 2013; Nagaoka et al. 2012). A majority of human aneuploidies originate from chromosome segregation errors in the female germline and increase dramatically with advancing maternal age (Chiang et al. 2010; Hodges et al. 2005; Lister et al. 2010; Yun et al. 2014a). This maternal-age effect reflects the deterioration of oocytes during the protracted arrest stage called dictyotene that is peculiar to oogenesis. Oocytes enter meiosis during fetal life, complete chromosome pairing and crossing over, and then arrest around birth. Resumption of meiosis and completion of the first division occurs only in mature oocytes that are about to be ovulated. Thus, an individual human oocyte can remain arrested in dictyotene for ∼11 to 50 years, i.e. puberty through menopause.

During the fetal stages of oocyte meiosis, chromosomes are replicated and sorted into homologous pairs, which then recombine to form inter-homolog crossovers (Hunter 2017; Reichman et al. 2017). In combination with cohesion between sister-chromatids, crossovers establish connections called chiasmata. When oocytes resume meiosis, chiasmata play an essential role in homolog disjunction, enabling homolog pairs to stably biorient on the meiosis-I spindle (MacLennan et al. 2015; Watanabe 2012). Homolog disjunction also requires that the two kinetochores of each pair of sister chromatids attach to spindle microtubules emanating from the same pole (mono-orientation) (Hauf et al. 2007; Hirose et al. 2011). At anaphase I, cohesion between the arms of sister chromatids is destroyed to resolve chiasmata and allow homolog disjunction. By contrast, cohesion between sister centromeres must be protected from destruction until anaphase II so that sister chromatids can biorient and disjoin on the meiosis II spindles (Holt and Jones 2009; Kudo et al. 2006).

It is now recognized that deterioration of sister-chromatid cohesion is a major underlying cause of the maternal-age effect (Chiang et al. 2010; Duncan et al. 2012; Lister et al. 2010; Yun et al. 2014b). Cohesin complexes establish cohesion during meiotic S-phase and early prophase I, but cannot be replenished thereafter l (Burkhardt et al. 2016; Revenkova et al. 2010; Severson and Meyer 2014; Tachibana-Konwalski et al. 2010). Thus, the depletion of cohesion in aging oocytes appears to be irreversible and impacts meiotic chromosome segregation in several ways. With deterioration of arm cohesin, chiasmata loosen causing them to migrate to more terminal positions (chiasma terminalisation), before eventually being lost, negatively impacting the ability of homologs to biorient and disjoin at meiosis I (Imai and Moriwaki 1982; Jagiello and Fang 1979; Kouznetsova et al. 2007; Lacefield and Murray 2007; Nagaoka et al. 2011; Sakuno et al. 2011). When centromeric cohesion is depleted prior to meiosis I, the normally tight association of sister kinetochores is weakened, increasing the risk of aberrant biorientation and equational segregation of sister chromatids at anaphase-I (Angell 1991; Angell et al. 1994; Shomper et al. 2014; Tachibana-Konwalski et al. 2013; Watanabe 2012). Even if sister kinetochores mono-orient and homologs disjoin during meiosis I, diminished centromere cohesion, including failure to protect centromere cohesion from destruction in anaphase I, can cause premature separation of sister centromeres (PSSC) and missegregation in meiosis II (Herbert et al. 2015).

In addition to maternal age, other extrinsic factors can contribute to oocyte segregation errors, including exposure to environmental toxins and being overweight or obese (Hunt et al. 2003; Luzzo et al. 2012; Wang et al. 2011). The effects of overweight and obesity on female reproduction have been studied for decades and are now generally accepted to have adverse impacts on pregnancy outcomes (Broughton and Moley 2017; Jungheim et al. 2010; Lashen et al. 2004; Selesniemi et al. 2011; Zhang et al. 2010). Weight gain and obesity are also generally associated with maternal aging (Goncalves et al. 2017; Pfannenberg et al. 2010). In both humans and mouse, either paternal or maternal obesity can impact embryo development and pregnancy outcome (Binder et al. 2012; Campbell et al. 2015; Finger et al. 2015; Fullston et al. 2015; McPherson et al. 2015), and even the health of offspring via transgenerational inheritance of epigenetic alterations (Ge et al. 2014; Saben et al. 2016; Watkins et al. 2018; Wu et al. 2015). The lower fertility of obese mothers is believed to derive from reduced oocyte quality because it can be ameliorated via *in vitro* fertilization with donor oocytes from normal weight individuals (Jungheim et al. 2013; Luke et al. 2011). However, other studies argue that obesity is also associated with lower uterine receptivity (Bellver et al. 2010; Bellver et al. 2013).

Mouse models have revealed that obesity can impact oocyte quality in several ways, including lipotoxicity, mitochondrial dysfunction, altered DNA methylation and increased apoptosis (Ge et al. 2014; Han et al. 2018; Jungheim et al. 2010; Saben et al. 2016; Wu et al. 2015; Wu et al. 2010; Zhao et al. 2017). Meiotic abnormalities associated with maternal obesity include defects in spindle morphology and chromosome misalignment (Boots et al. 2016; Han et al. 2017; Luzzo et al. 2012). In later life, obesity can develop into diabetes, and maternal diabetes in mouse has been associated with increased rates of PSSC and aneuploidy (Cheng et al. 2011; Wang et al. 2009).

In the current study, we explore the associations between maternal obesity, chromosome abnormalities and age-dependent exhaustion of cohesion. Our data suggest that obesity exacerbates the maternal-age effect by accelerating cohesin deterioration.

## RESULTS

A well-established mouse model of obesity induced by high-fat diet (HFD) (Ge et al. 2014) was employed to investigate the effects of maternal obesity on oocyte quality, focusing on events at the chromosome level. Animals at three and six-months old, fed either HFD or a control diet (CD), defined our four experimental cohorts. Consistent with previous studies (Ge et al. 2014), at three months old, body weights of HFD-fed animals were 44.5% higher than those of controls (**Fig. 1A**; 29.5 ± 6.2 g versus 20.4 ± 1.6 g in CD controls, *P*=0.0015, *t*-test); and by six months HFD animals were 68.8% heavier than controls (**Fig. 1A;** 38.5 ± 4.4 g versus 22.8 ± 2.7 g in CD controls, *P*<0.0001, *t*-test).

**Figure 1:**
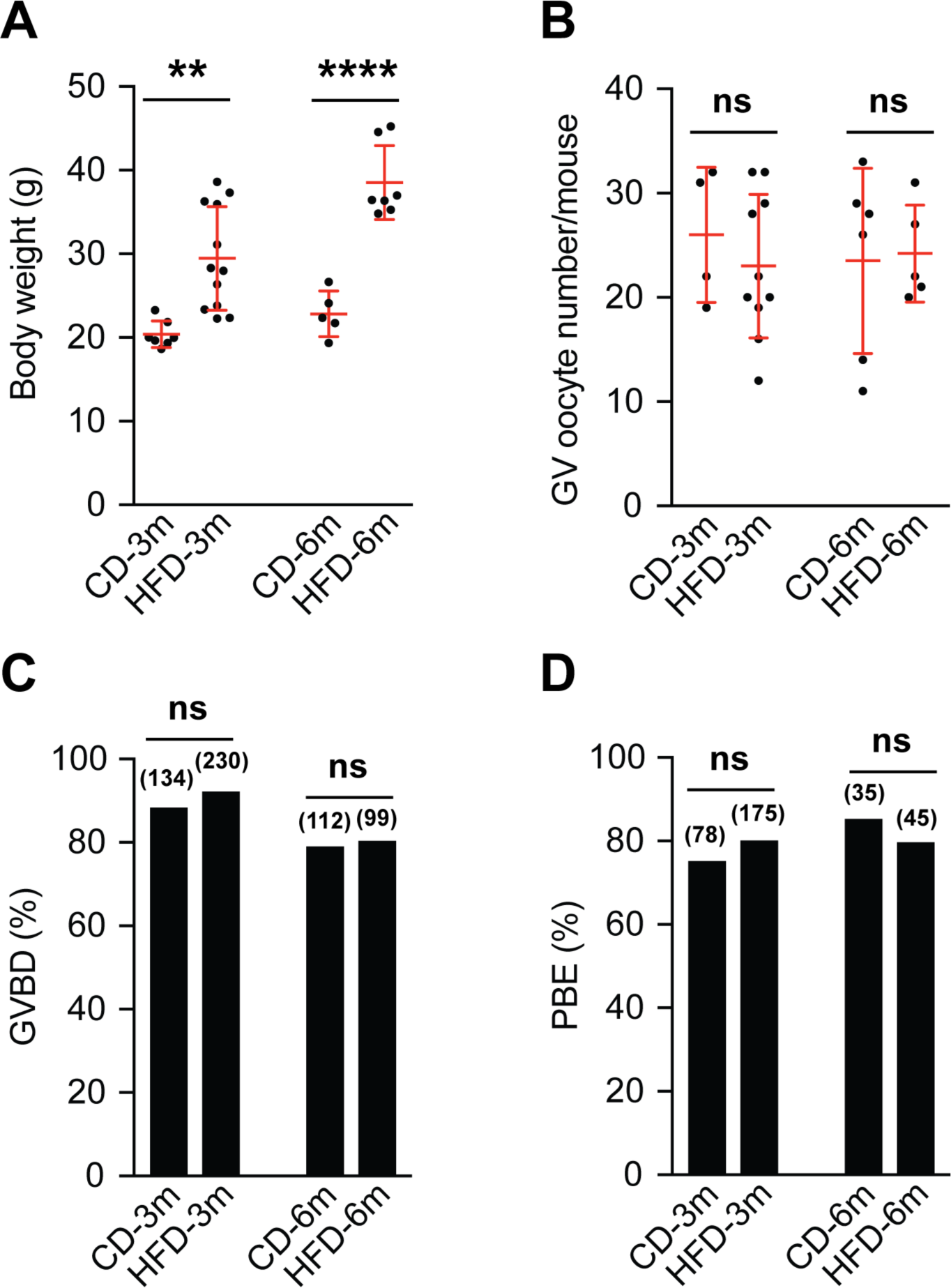
Obesity does not affect oocyte maturation. **(A)** Body weights of 3-month (3m) and 6-month (6m) old experimental animals (mean ± s.d.). **(B)** Numbers of GV oocytes retrieved per animal (mean ± s.d.). (**C** and **D**) Proportions of oocytes undergoing GVBD (**C**) and PBE (**D**). Numbers of oocytes analysed are indicated in parentheses. Data were analysed using the Student’s t-test (A and B) or Fisher’s exact test (C and D). **, *P*<0.01; ****, *P*<0.0001; ns, not significant.

### Obesity does not affect the efficiency of oocyte maturation

Effects of maternal obesity on oocyte growth and maturation were assessed by: (i) quantifying the numbers of fully grown germinal-vesicle (GV) oocytes retrieved from the ovaries of aged-matched HFD and CD animals; (ii) the efficiency of meiotic resumption when oocytes were cultured *in vitro*, as assessed by GV breakdown (GVBD); and (iii) the efficiency of the meiosis-I division scored by first polar body extrusion (PBE; **Fig. 1B,C,D**).

Surprisingly, there were no significant differences between the numbers of GV-stage oocytes retrieved from any of the four study cohorts. Though variation among individual animals from each group was large, the mean values from all groups were similar, ranging between 23 and 26 GV oocytes per animal (**Fig. 1B**). Contradicting previous analysis (Hou et al. 2016), we detected no effects of maternal obesity on oocyte maturation when cultured *in vitro.* The efficiencies of both GVBD and PBE were indistinguishable for oocytes from HFD and CD animals (**Fig. 1C, D**).

### Obesity is associated with chromosome abnormalities in both metaphase I and II of oocyte meiosis

Metaphase chromosome spreads of *in vitro* cultured oocytes were used to assess abnormalities in both MI and MII (**Fig. 2A**). Abnormalities detectable in MI include unconnected univalents caused by either failed crossing over during fetal oogenesis, or premature loss of cohesion between chromosome arms. Univalents are prone to missegregation at anaphase I, either mono-orienting on the MI spindle and disjoining as intact pairs of chromatids (homolog non-disjunction); or becoming bi-oriented and undergoing equational segregation. Consequently, abnormalities detectable in metaphase-II arrested oocytes include numerical abnormalities (aneuploidy) and free chromatids (premature separation of sister centromeres, PSSC).

**Figure 2:**
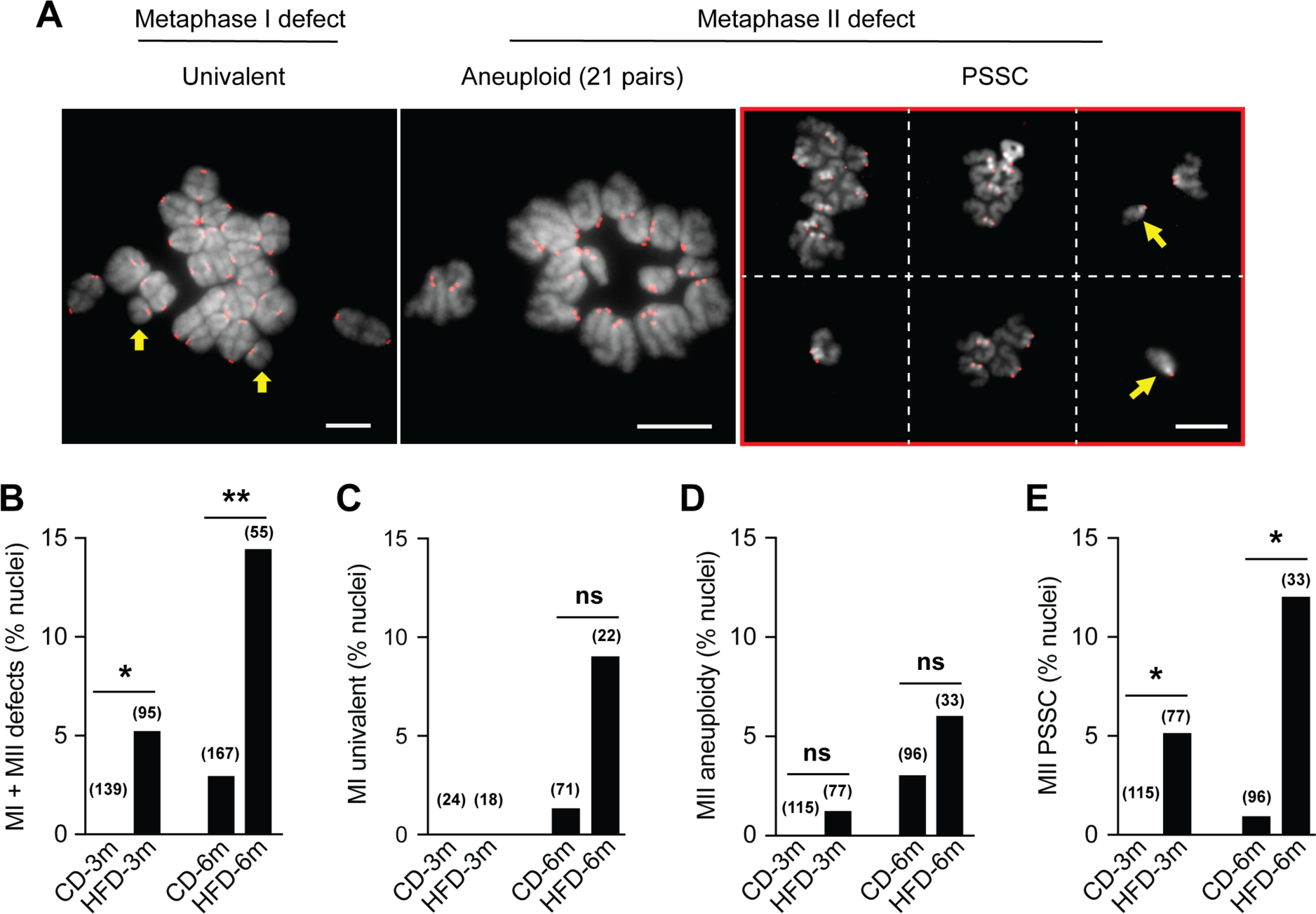
Chromosome abnormalities are increased in oocytes from obese mice. **(A)** Representative images of chromosome abnormalities detected in metaphase-I and metaphase-II chromosome spreads. Left to right: metaphase-I oocyte with 19 bivalents and 1 pair of univalents (yellow arrows); aneuploid metaphase-II oocyte with 21 dyads; euploid metaphase-II oocyte with PSSC (yellow arrows highlight free chromatids; images in the red rectangle are from the same cell, but represent different fields of view). Kinetochores (CREST) are red, chromosomes (DAPI) are grey. Scale bars, 10 µm. (**B–E)** Quantification of **(B)** all oocytes with chromosome abnormalities; (**C**) metaphase-I oocytes with univalents; (**D**) aneuploid metaphase-II oocytes; (**E**) metaphase-II oocytes with PSSC. Numbers of oocytes analysed are indicated in parentheses. *, *P*<0.05; **, *P*<0.01; ns, not significant; Fisher’s exact test.

Both three and six month HFD obesity cohorts showed significantly elevated levels of oocyte chromosome abnormalities relative to CD controls (**Fig. 2B**). 9.1% (2/22) of metaphase-I oocytes from six-month old HFD animals contained univalents compared to 1.4% (1/71) of CD controls, but this difference was not statistically significant at the <5% level (*P*=0.14, fisher’s exact test; **Fig. 2C**). Univalents were not detected in three-month old animals, either HFD or CD.

Similarly, in metaphase-II arrested oocytes, aneuploidy was detected at low levels but differences between HFD and CD cohorts were not statistically significant (**Fig. 2D**). However, significant increases in metaphase II oocytes with PSSC were observed in HFD animals (**Fig. 2E**). In three month-old animals, 5.2% (4/77) of metaphase II eggs contained single chromatids, which were never observed in the age-matched CD group (0/115; *P*=0.02, Fisher’s exact test). PSSC was even more frequent in six month-old HFD animals, with 12.1% (4/33) oocytes containing single chromatids compared to just 1.0% (1/96) in age-matched controls (**Fig. 2E**). Notably, all nuclei with PSSC contained two free chromatids but were otherwise euploid (n=40 chromatids). These abnormalities most likely derive from normal homolog disjunction in MI, accompanied by complete loss of centromere cohesion between a single chromatid pair (Herbert et al. 2015).

### Chiasma terminalization is increased in the oocytes of obese animals

Deterioration of sister-chromatid cohesion can lead to univalents in metaphase I, and aneuploidy and PSSC in metaphase II. To investigate whether cohesin deterioration contributes to obesity-related increases in chromosome abnormalities, we analysed the distributions of chiasmata in metaphase-I chromosome spreads (**Fig. 3**). We scored three classes of chiasmata based on the numbers and positions of chiasmata: (i) terminal, with a single end-to-end connection; (ii) interstitial, with a single connection and visible cross-shaped morphology; and (ii) multiple, with two or more connections (**Fig. 3A**).

**Figure 3:**
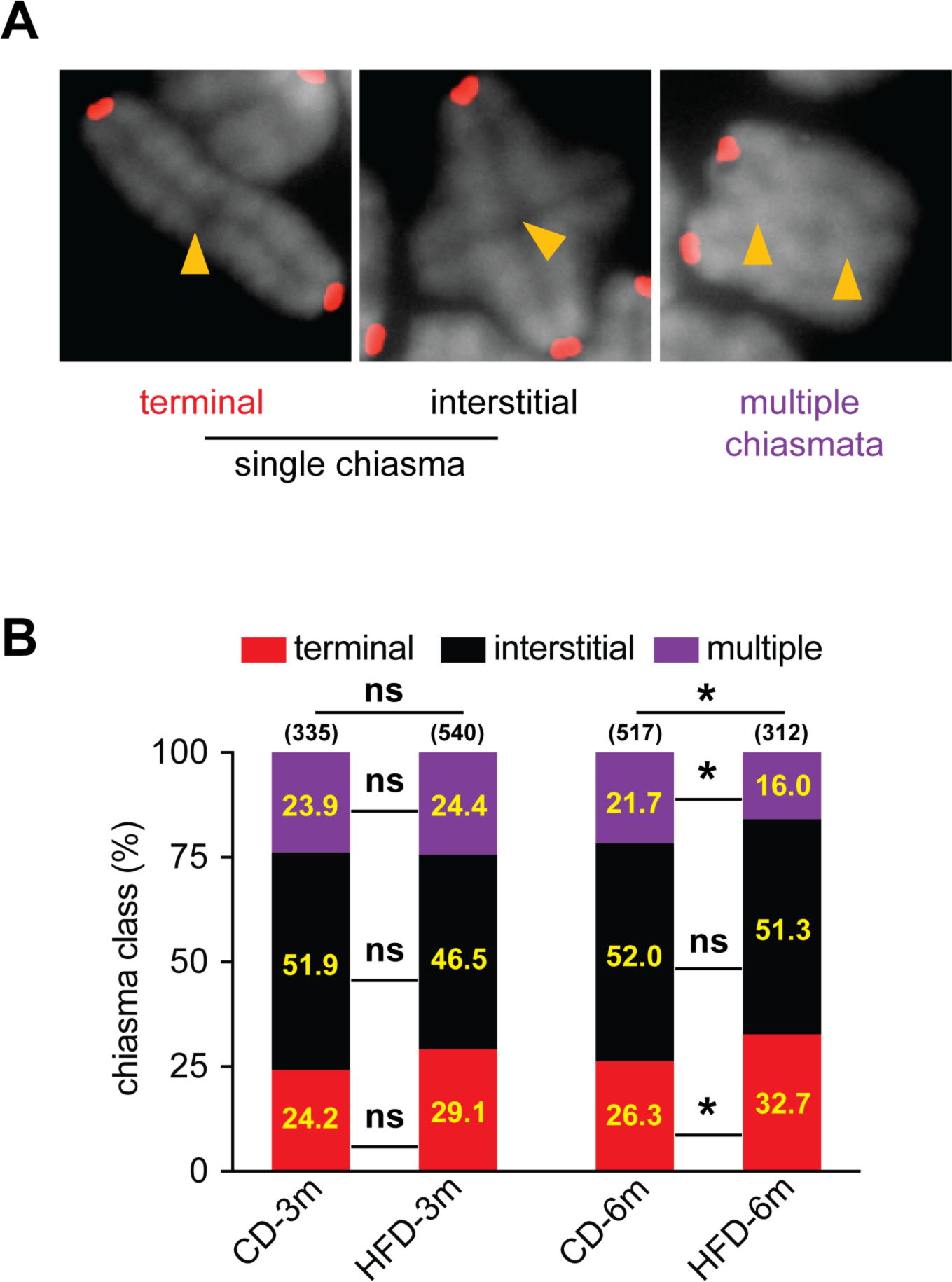
Chiasma terminalisation is increased in metaphase I oocytes of obese mice. (**A**) Representative images of bivalents with terminal, interstitial and multiple chiasmata. Chiasma positions are highlighted with yellow carets. (**B**) Chiasma distributions. Numbers in the sub-bars show the percentage of each chiasma class. Numbers of bivalents analysed are indicated in parentheses. *, *P*<0.05; ns, not significant; *G*-test for distributions, Chi-square test for pairwise comparisons.

In oocytes from six-month old animals, the distributions of chiasmata classes were significantly different between the HFD and CD cohorts (*P*<0.05; *G*-test). HFD oocytes had elevated levels of terminal chiasmata (32.7% versus 26.3% in CD oocytes, *P*<0.05, Chi-square test) and corresponding reductions in bivalents with multiple chiasmata (16.0% versus 21.7% in CD oocytes, *P*<0.05, Chi-square test; **Fig. 3B**). By comparison, distributions of chiasmata classes did not differ between three-month old HFD and CD animals. Thus, maternal obesity appears to accelerate the age-dependent deterioration of cohesin.

### Inter-kinetochore distances are increased in metaphase II oocytes of obese animals

Increased terminalization of chiasmata in HFD oocytes of 6-month old animals indicates accelerated deterioration of cohesion between chromosome arms. However, the most prominent chromosomal abnormality observed in HFD oocytes, PSSC, occurs when centromeric cohesion is diminished (Chiang et al. 2010; Yun et al. 2014b).

To assess whether centromere cohesion is weakened in obese mice, inter-kinetochore distances (IKDs) were measured in metaphase-II arrested oocytes (**Fig. 4A**). Analysis of the average IKDs per nucleus showed a trend towards slightly larger IKDs in HFD oocytes from three-month old animals (1.11 ± 0.22 µm versus 1.04 ± 0.19 µm in CD controls; **Fig. 4B**), but the difference was not significant (*P*=0.24, Mann-Whitney test). However, the rank distributions of IKDs for individual sister-chromatid pairs indicated significantly larger IKDs in three-month old HFD oocytes (*P=*0.004, Mann-Whitney test; **Fig. 4C**). At six months, significant increases in IKDs were readily detected in HFD oocytes, both by analysis of average IKDs per nucleus (1.31 ± 0.22 µm versus 1.10 ± 0.27 µm in CD controls, *P=*0.01, *t-* test; **Fig. 4B**) and by the rank distributions (*P<*0.0001, Mann-Whitney test; **Fig. 4D**). Thus, HFD-induced obesity is also associated with accelerated weakening of centromeric cohesion.

**Figure 4:**
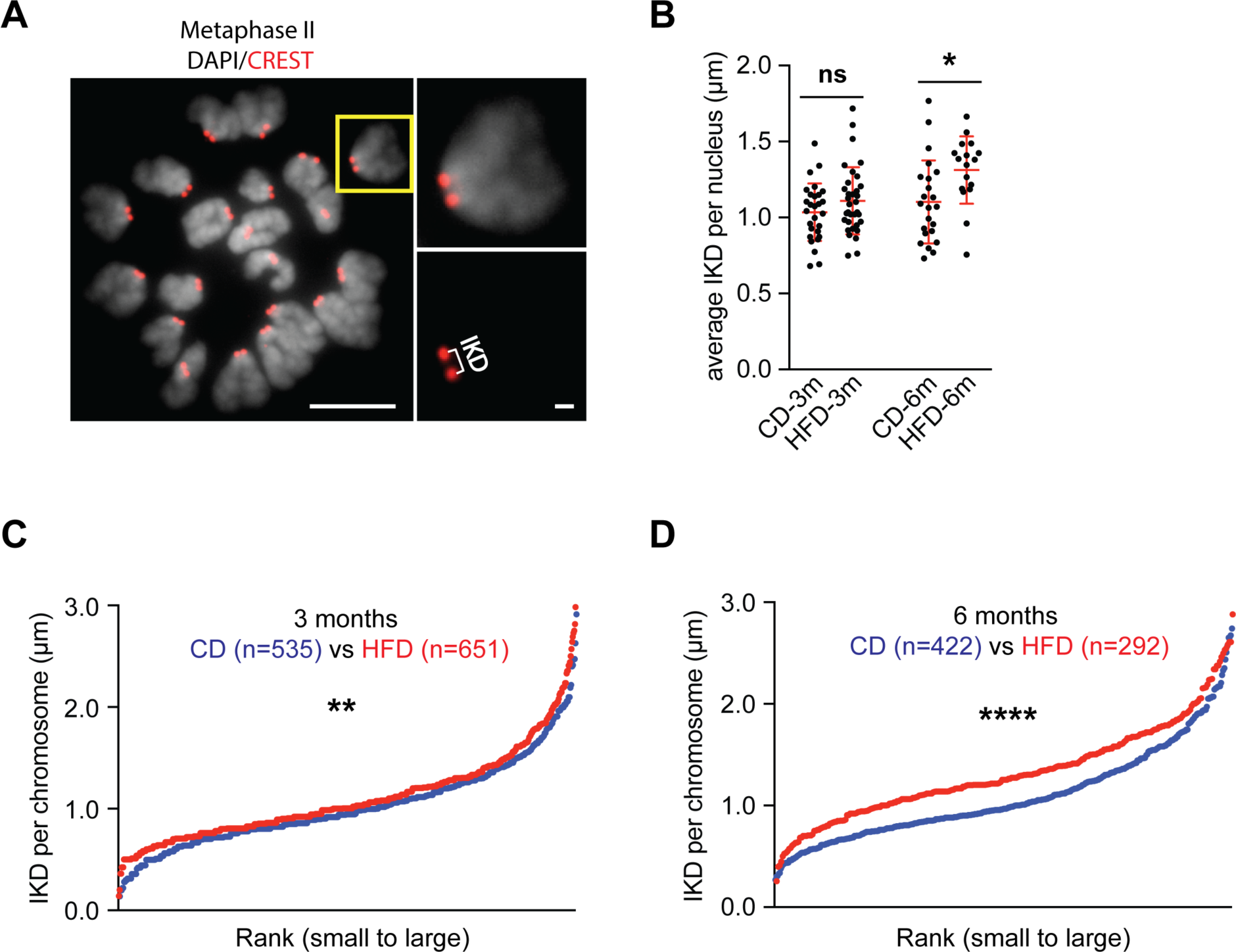
Inter-kinetochore distances are increased in metaphase II oocytes from obese mice. (**A**) Representative chromosome spread of a metaphase II oocyte. A single dyad is enlarged to highlight the IKD measurement. Kinetochores (CREST) are red, chromosomes (DAPI) are grey. Scale bars represent 10 µm (nucleus) or 1 µm (dyad). (**B**) Average IKDs per nucleus. Bars show means ± s.d. of all nuclei. (**C, D**) Rank distributions of the IKDs of individual dyads. *, *P*<0.05; **, *P*<0.01; ****, *P*<0.0001; ns, not significant; Student’s *t*-test (6m group in B); Mann-Whitney test (3m group in B, C, D).

## DISCUSSION

In our study, maternal obesity did not affect the numbers of fully-grown GV oocytes produced, or their ability to mature *in vitro* (**Fig. 1B-D**). Similarly, the data of Ata *et al.* showed that Body Mass Index in humans did not affect oocyte yield or *in vitro* maturation (Ata et al. 2013). However, conflicting data from human *in vitro* fertilization (IVF) studies have indicated that maternal obesity does negatively impact oocyte yield and maturation (Zhang et al. 2010; Zhang et al. 2015a). IVF routinely involves long-term administration of gonadotropin to stimulate follicle growth and increase numbers of mature eggs. It is important to note that obese women, who have a higher risk of infertility, show a reduced response to gonadotropin (Ozekinci et al. 2015; Zhang et al. 2010). Thus, the reduced yield and maturation efficiency of eggs from obese women in an IVF setting could be caused primarily by their decreased response to gonadotropin. Consistently, minimizing ovary stimulation of obese women can negate these effects (Zhang et al. 2015a) Importantly, in our mouse study, oocyte growth was not primed with gonadotropin and obese mice yielded normal numbers of healthy oocytes capable of reentering meiosis and completing MI with normal efficiency.

Although obese mice produced normal numbers of oocytes, their quality was reduced at the chromosome level, with enhanced deterioration of cohesion leading to chiasma terminalization, increased IKDs and PSSC. These data complement a previous study (Luzzo et al. 2012), which showed an association between maternal obesity, MI spindle defects and chromosome misalignment in mouse oocytes. Importantly, obesity mimics and thereby exacerbates the deterioration of cohesion seen during maternal aging. This observation suggests that obesity and maternal aging could diminish cohesion via similar mechanisms. A possible unifying mechanism of cohesion deterioration is oxidative damage. HFD-induced obesity leads to increased oxidative stress in mouse oocytes (Jia et al. 2018; Zhang et al. 2015b); and Perkins et al. recently showed that genetically-enhancing oxidative stress increases cohesion loss and segregation errors in *Drosophila* oocytes (Perkins et al. 2016). Oxidative stress could damage the cohesins themselves as well as factors directly or indirectly involved in their stability.

PSSC events detected in HFD oocytes, comprising two free chromatids in an otherwise euploid nucleus, are consistent with normal homolog disjunction in MI, followed by dissociation of a single pair of chromatids. Such events could be the consequence of a general deterioration of cohesion during dictyotene, or a more specific but not exclusive failure to protect centromeric cohesion from destruction in anaphase I. The first possibility predicts that HFD oocytes will have increased IKDs in metaphase I, while the latter predicts that only metaphase II oocytes will show significantly increased IKDs and PSSC. Favoring a model in which HFD-induced obesity leads to inefficient protection of centromeric cohesion, we did not detect increases in average IKDs per nucleus in metaphase-I oocytes (0.53 ± 0.08 µm in HFD-6m versus 0.56 ± 0.08 µm in CD-6m controls, *P=*0.31, *t*-test).

At the onset of anaphase I, the separase protease is released from inhibition and cleaves specifically phosphorylated cohesin to allow homolog disjunction (Holt and Jones 2009). Centromere-associated cohesin is protected from cleavage by the PP2A phosphatase, which is recruited to centromeres by the Shugoshin protein, Sgo2 (Kitajima et al. 2006). Interestingly, Sgo2 is also reduced with advancing maternal age (Lister et al. 2010; Yun et al. 2014b). Given our observations, we propose that obesity may also lead to diminished Sgo2 levels. In summary, our data provide new insight into the complex effects of obesity on female reproduction, revealing enhanced cohesin deterioration that mimics and compounds the maternal-age effect.

## METHODS

### Animals and Diet

Original C57BL/6J mice were purchased from the Jackson Lab. Animals were maintained and used for experimentation according to the guidelines of the Institutional Animal Care and Use Committees of the University of California, Davis. Weanling females (21 days post-partum) were housed 3 or 4 per cage and fed *ad lib*. The mice were fed either a high-fat diet (HFD; Research Diets Inc.; D12492 with 60% kcal% fat) or a standard rodent chow control diet (CD) for 3 or 6 months.

### Oocyte collection and *in vitro* maturation

Germinal-vesicle stage oocytes were collected from ovaries of experimental animals without prior hormonal stimulation. Only oocytes with integral cumulus cell layers were utilized. Oocytes were collected in M2 medium under mineral oil at 37°C, and cultured for *in vitro* maturation after mechanically removing surrounding cumulus cells for observation of Germinal Vesicle Breakdown (GVBD) and Polar Body Extrusion (PBE). GVBD was scored after 3 hours of culture. Only oocytes that underwent GVBD within 3 hours were used to quantify PBE after a further 12 hours of incubation.

### Chromosome spreads of metaphase I and II oocytes

Metaphase I and II oocytes were acquired after 7 hours and 15 hours of culture in M2 medium at 37°C. Metaphase chromosome spreads were performed as previously described (Qiao et al. 2018). Briefly, Acid Tyrode’s solution (Sigma-Aldrich) was applied to remove zona pellucida, and zona-free oocytes were spread in 1% paraformaldehyde with 0.15% Triton-X-100 and 3 Mm DTT in H_2_O. To improve the accuracy of chromosome counting, single oocytes were spread in each well of a 12-well slide (Electron Microscopy Science). Spreads were air dried at room temperature overnight.

### Immunofluorescence and data analysis

Metaphase I and II oocyte spreads were blocked at room temperature for 1-2 hours, followed by overnight incubation with primary antibody – human anti-centromere antibodies (ACA or CREST; HCT-0100 ImmunoVision, 1:1000). Goat anti-human 555 secondary antibodies (Molecular Probes, 1:1000) were incubated for 1hour at room temperature. Chromosomes were stained with 4′,6-diamidino-2-phenylindole (DAPI). Images were acquired using a Zeiss AxioPlan II microscope equipped with a 63X Plan Apochromat 1.4 objective, EXFO X-Cite metal halide light source with Hamamatsu ORCA-ER CCD camera. Image processing and analysis were performed using Volocity (Perkin Elmer) and ImageJ (NIH) software.

### Statistical analysis

Dichotomous data were analysed using Fisher’s exact, Chi-squared or *G* tests. All other mean analyses were performed using either the Student’s t or Mann–Whitney tests. Data were processed using GraphPad Prism, with a significance threshold of *P* < 0.05. In all comparisons, at least two mice (littermates or age-matched) were examined from at least two independent experiments.

## Acknowledgments

We thank Richard Shultz and members of the Hunter Lab for support and discussions. N.H. is an Investigator of the Howard Hughes Medical Institute, which also supported this study.

